# Sensory and developmental phenotyping of *C. elegans* parses autism associated genes into behavioural classifications

**DOI:** 10.64898/2026.03.27.714775

**Authors:** James W LaVilliers, Eleonora M Pieroni, Farah Al Khawaja, Katrin Deinhardt, Vincent M O’Connor, James C Dillon

## Abstract

Differences in sensory processing and development are traits described by the diagnostic criteria for autism spectrum disorder (ASD). The inclusion of sensory processing has been accompanied by increased investigation of this criterion to better understand how autism risk genes bring about changes in sensory biology. The contribution of epigenetic modifiers to sensory and developmental phenotypes is of particular interest, given they represent genetically encoded signalling cascades that regulate the environments’ ability to modulate gene expression. ∼50% of the highest confidence genes ranked on the Simons Foundation Autism Research Institute (SFARI) can be classified as epigenetic modifiers, further highlighting the contribution of aberrant epigenetic function in shaping the landscape of ASD. Here, we utilise the model organism *C. elegans* to investigate epigenetic modifiers, their role in sensory and developmental biology and behavioural outcomes.

*C. elegans* provides robust readouts for both developmental and sensory biology allowing phenotypic signatures of ASD to be systematically investigated. 52 epigenetic modifiers (65 strains) were selected for study in *C. elegans* based on gene function, presence of an orthologue in *C. elegans* and the availability of viable putative null strains. This identified significant changes to reproduction, gross development and sensory processing across the range of epigenetic modifiers. We identified subgroups of mutant strains with either impaired sensation or development, or both. This study provides a link between the ASD phenotypes of sensory impairment and developmental delay and suggests sensory testing in human cohorts can help to further refine sub-categorisation of heterogenous ASD cohorts.

## Introduction

Autism spectrum disorder (ASD) is a spectrum of conditions clinically diagnosed by two behavioural features encompassing social communication/interaction and restricted/repetitive patterns of behaviour (American Psychiatric Association, 2013). These behavioural traits are diagnosed during early development and then persist with varying levels of severity (American Psychiatric Association, 2013). In addition, perturbation of sensory processing in ASD cohorts has been observed across multiple modalities (Balasco et al., 2020) and considered an important diagnostic trait due to their high incidence (Kim et al., 2025). However, this has not been systematically coupled to other clinical measurements that are used for ASD diagnosis. Sensory testing ranges from parent’s self-reporting behaviours (Balasco et al., 2020) to matched cohort sensory testing in laboratory conditions (Balasco et al., 2020).

The understanding of the genetic architecture of ASD is improving, with the increasing quantitative nature of these investigations allowing genotype and associated traits to be grouped to inform on underlying mechanisms (Leyhausen et al., 2025; Litman et al., 2025). With heritability rates ranging from 75%-95% (Colvert et al., 2015), the contribution of genetic variation to the aetiology of ASD is unequivocal but genetics alone is unable to provide a wholistic explanation of the disorder. The evidence for a genetic contribution is rooted in the abundance of genes associated with ASD discovered through genome wide association studies (GWAS) (Grove et al., 2019), whole exome sequencing (WES) (Yu et al., 2013) and twin studies (Tick et al., 2016). The weight of the evidence supporting an association with identified genetic risk factors is catalogued and ranked on the Simons Foundation Autism Research Initiative (SFARI) gene database with over 1200 (Jan 2026) (Abrahams et al., 2013) genes implicated in the aetiology of the disorder.

Genetic variation does not account for all ASD cases and accumulated evidence supports a role for disrupted epigenetic programming of nervous system function and development (LaSalle., 2023). Epigenetic modifiers are found abundantly in the SFARI gene database (Abrahams et al., 2013). This makes these genes particularly interesting to study in the aetiology of ASD because they co-ordinate molecular programmes in response to environmental changes required for context dependent outputs (LaSalle., 2023). Hence, these genes can inform on how the environment impacts neuronal function and mechanisms underpinning ASD. (Loke, Hannan and Craig., 2015).

Understanding the intersection between genetics and the environment will enable classifications that parse the ASD diagnosis into biologically distinct sub-cohorts. Indeed, a recent study has validated that ASD cohorts can be statistically parsed by combining phenotypic presentation of clinical symptoms including developmental milestones, repetitive behaviours, social communication and child behaviour and associated genetic signature (Litman et al., 2025). The four key groups were sub-categorised as moderate challenges, broadly affected, social and/or behavioural and mixed ASD with developmental disorder (DD). Findings from another study seeking to investigate possible subgroupings of ASD cohorts identified three sub-categories of autistic individuals (Leyhausen et al., 2025). These subgroups were established based on distinct neuroanatomical phenotypes and transcriptomic profiles (Leyhausen et al., 2025).

Interestingly the aforementioned sub-categorisation models are built without clear use of the association of neuroatypical sensory sensitivity to environmental cues. Hyper or hyporeactivity to sensory input is commonly observed in individuals with ASD, it has been postulated to originate in altered neural pathways (Palau-Baduell et al., 2012), atypical sensory modulation (Giacardy et al., 2018) and sensory gating dysfunction (Chien et al., 2019). A definitive explanation for sensory perturbations has yet to be uncovered (Balasco et al., 2020) but has excellent potential to refine the classification of emerging subgroupings (Table S1).

In this study we have used *C. elegans* to conduct a phenotypic classification of ASD associated gene modifiers that function in epigenetic regulation. As a model system, *C. elegans* has the advantage of genetic tractability combined with a diverse range of quantitative assays for interrogating behavioural responses to external chemosensory cues. Furthermore, understanding changes to chemosensory processing can inform upon developmental outcomes as sensory impairments have been shown to cause developmental delays in *C. elegans* (Fujiwara et al., 2002; Rose et al., 2005).

This work builds upon previous research leveraging *C. elegans* to understand genotype/phenotype interactions in specific areas of ASD aetiology. These include morphological/developmental changes (Aguirre-Chen et al., 2020; McDiarmid et al., 2019; Wong et al., 2021; Zoga et al., 2025), social (Rawsthorne et al., 2021, Cowen et al., 2024), sensory (Schmeisser et al., 2017) and learning behavioural readouts (McDiarmid et al., 2019).

Here we have quantified *C. elegans* development and chemosensory responses across gene expression regulators associated with a high risk of ASD. The pipeline readily allowed scoring of the integrity of sensory responses and development, allowing strains to be organised into distinct phenotypic groupings. The groupings showed either “sensory” or “developmental” or combined “development, reproduction and sensory” scores relative to the wild-type strain (N2). This phenotypic grouping, built upon sensory and developmental analysis further supports the sub-categorisation of ASD risk associated genes by phenotypic criteria. The analysis described here, highlights the value of sensory measures in establishing biologically distinct sub-classifications of ASD.

## Materials and Methods

### *C. elegans* strains and culturing

The N2 (Wild-type, Bristol Strain) was obtained from the *Caenorhabiditis* Genetics Center (CGC, University of Minnesota, USA.) The remaining strains were obtained from the CGC and National Bioresource Project and made by S. Mitani (NBRP) (Yamazaki et al., 2010) (Table S2). All strains were cultured at 20°C on standard nematode growth medium (NGM). The worms were maintained on *Escherichia coli* strain OP50 (Brenner, 1974).

### Bioinformatics

Category 1 and category S genes listed on SFARI (Accessed, Jan 2026, Ver 2025 Q4, https://gene.sfari.org/), were used for candidate gene selection. Category 1 genes are classified as high confidence with >three de novo gene disrupting mutations having been reported. Category S genes are classified based on the consistent link to additional characteristics not required for an ASD diagnosis. Gene expression modifiers were identified by review of the literature using the descriptor ‘gene expression modifying activity’ as the criteria to functionally annotate the genes selected from the SFARI database. The key GO descriptors were ‘transcriptional regulation’, ‘translational regulation’ and ‘regulation of gene expression’. Ortholist2 (http://ortholist.shaye-lab.org/) (Kim et al., 2018) and STRING (https://string-db.org/) (Szklarczyk et al., 2023) were used to determine whether *C. elegans* has an orthologue of the gene expression modifiers identified in the previous step. Viable mutants were identified using Wormbase (Version: WS297; https://wormbase.org/species/c_elegans#401--10) which contains JBrowse2 for strain identification and reported phenotypic information on available strains such as the effect of the mutation and only selecting those with a high variant effect predictor score (VEP). This enabled exclusion criteria such as sterility, lethality and severe motor defects to be applied to each candidate gene strain. Additional information on the expression of the human orthologue of the candidate genes was collated using Bgee (https://www.bgee.org/) (Bastian et al., 2021) and *C. elegans* transcript number data using CeNGEN (https://cengen.shinyapps.io/CengenApp/) (Taylor et al., 2021).

Sequence alignments of amino acids for identity and similarity comparison of orthologues was performed with EMBOSS Needle (https://www.ebi.ac.uk/Tools/psa/emboss_needle/) (Madeira et al., 2024). Where multiple orthologues were present, the sequence alignments were used to identify the closest matching orthologue for use within this study.

### Egg Laying and Development Analysis

A synchronised population of hermaphrodite worms was generated by allowing 3 L4+1-day (reproductive young adults) worms to lay eggs for 3 hours before removal. Worms were visually inspected for the presence of eggs prior to picking onto the assay plate and only worms containing eggs were used. This was then converted to an egg laying rate (number of eggs/number of worms/time). Developmental analysis was performed on the same plates used for egg laying and recorded by allowing the eggs to develop at 20°C. At set time points, 0 hours, 20 hours, 25 hours, 43 hours, 48 hours and 67 hours the plates were observed under a stereo microscope (Nikon SMZ800N) to count the number and developmental stage of the worms. These stages are defined as either eggs, L1 or L2 for early development and L3, L4 and young adults for late development as previously described (Thummel, 2001). All strains were compared to N2 and the distribution and variation of the distinct stages used to determine the developmental timings. Additionally, *che-3 (e1124)* (lacking all sensory amphids and phasmids) was included to benchmark the effects of sensory deficits on development in *C. elegans*. Strains were compared to N2 and rank ordered for developmental delays. A correlation between early and late developmental rank orders for each strain was then performed. Experimenters were blind to the strains being investigated during the assay and statistical analysis.

### Confocal Microscopy and DiI Labelling

A synchronised population of L4+1-day (72 hours old) worms were washed once with 1 ml M9 before being resuspended in 1 ml DiI in M9 (2 mg/ml Invitrogen, Catalogue: D282) and left on a shaker for 3 hours at room temperature. Worms were washed 3 times post staining and re-plated onto an OP50 seeded NGM plate for 15 minutes prior to imaging. Individual worms were picked into a single drop of 10 mM sodium azide (CAS 26628-22-8, Sigma-Aldrich) solution on a 1% agar pad, on a microscope slide to paralyse them for imaging (Schultz and Gumienny, 2012).

Worms were then imaged using a Leica DMI8-CS microscope with excitatory laser DPSS 561 nm and collection of emitted light at 565 nm – 660 nm. Following staining, sensory amphids ASK, ADL, ASI, AWB, ASH and ASJ and phasmids PHA and PHB were identified using their anatomical location and morphology (Andrews et al., 2025). The above protocol successfully stained the 6 amphid neurons and 2 phasmid neurons, listed previously, in all N2 worms. N2 staining was run in parallel to the mutant strains and used as a reference to compare the number and gross morphology of the sensory neurons in the mutant strains. Strains were deemed to be significantly different if at least one amphid neuron or phasmid neuron did not label. Experimenters were blind to the strains being investigated during the assay and image analysis.

### Quadrant Assay

The arena consisted of a 6 cm petri dish with 13 ml NGM. On the plastic base of the dish the plate was split into 4 equal sized quadrants. A dot was placed in each quadrant 2 cm from the intersect point of the quadrant lines. An immobile worm zone was also outlined at 0.5 cm from the intersect point in all directions. The quadrants were assigned as alternating test and control zones. On the day of the assay, 1 µl 0.5 M sodium azide was pipetted on the aforementioned dot in each quadrant first and allowed to dry. This was followed by either 1 µl M9 in the control quadrants or 1 µl 1% v/v diacetyl (Sigma-Aldrich, Catalogue: 8.03528) in M9. A synchronised population of worms was generated by allowing 7 L4+1-day worms to lay eggs for 4 hours, washed twice using 1ml M9 solution before being transferred to the assay plate with a glass capillary onto the intersect point of the quadrant in <15 µl M9. 50-100 worms were transferred to the assay plate. The assay plates were then left for 50 minutes and scored based on the number of worms in each quadrant. A chemotaxis coefficient was obtained by taking the differential of worms on the test vs control quadrants divided by the total number of worms in the assay. Worms still within the 0.5 cm radius immobile zone were counted as non-moving and the percentage of non-moving worms per mutant was recorded at the end of the assay and compared to N2. Experimenters were blind to the strains being investigated during the assay and statistical analysis.

### Droplet Avoidance Assay

We adapted the classical acute aversion assay first described by Hilliard et al., 2002; Hilliard et al., 2004. 9 cm plates were poured with NGM (20 ml) 3 days prior to the assay. 7 L4+1day old worms were picked onto an unseeded NGM plate and allowed to roam for 30 seconds to remove residual bacteria. These worms were then transferred onto a fresh unseeded 9 cm NGM plate and left for at least 20 minutes to allow the transition from local area search to escape behaviour (Gray et al., 2005).

One drop (<20 μl) of the indicated aversive cue was placed on the trajectory of a forward moving worm. Soluble aversive cues used in this study included copper sulphate (CuSO_4_, 30 mM), M9 (pH 3.0), and fructose (4M) in water and were controlled against M9 (pH 7.0). The volatile aversive cue was 30% v/v 1-octanol in ethanol and presented on a platinum wire in the trajectory of a forward moving worm.

The performance of the selected mutant strains was recorded and compared to the N2 and the sensory mutant *che-3 (e1124*) (sensory deficient control). *che-3 (e1124)* lacks all sensory amphids and phasmids (Wicks et al., 2000). Video recordings were made of the worm’s response to the sensory cue and analysed *a posteriori*. For soluble aversive cues (water soluble cues), the total number of reversals initiated within 5 seconds of compound exposure was counted. For the volatile aversive cue 1-octanol (30% v/v), the latency from cue introduction to initiating a reversal was recorded. Experimenters were blind to the strains being investigated during the assay and video analysis.

### Significance Heatmap and phenotypic criteria

Data from statistical tests of each ASD associated mutant in all assays conducted were compiled. For each strain and condition (except DiI labelling), statistically determined p values from post-hoc multiple comparison testswere compiled.

Developmental delay was defined as deviating significantly from N2 in either early or late development. Sensory impairment was defined as deviating significantly from N2 in either attractive or aversive (soluble or volatile) sensory processing. Severe sensory impairment was defined as significantly deviating from N2 in both attractive and aversive (soluble or volatile) sensory processing.

### Statistical Analysis

Statistical analysis was performed using GraphPad Prism 10.4.2. All data expressed as mean ±SEM. Statistical tests and post-hoc analysis are outlined in figure legends. Significance defined at p<0.05. Briefly, egg laying, development, attractive and aversive chemosensory processing were all analysed using a one-way ANOVA with Dunnett’s multiple comparison test.

## Results

### A bioinformatic pipeline for selection of ASD associated gene expression modifiers: comparative phenotypic analysis in *C. elegans*

The SFARI database was used and those listed as category 1 genes were selected for the current pipeline. Our intention was to bias the investigation to select for genes that encode modulators of gene expression. Of the total 232 genes classified as category 1, 111 (48%), met the indicated definition of being gene expression modifiers (see methods). Orthologues were identified using Ortholist 2 (Kim et al., 2018) and STRING (Szklarczyk et al., 2023). Further criteria refined the candidate gene pool to those with a *C. elegans* orthologue (Fig 1A, 76 genes).

**Fig 1.**
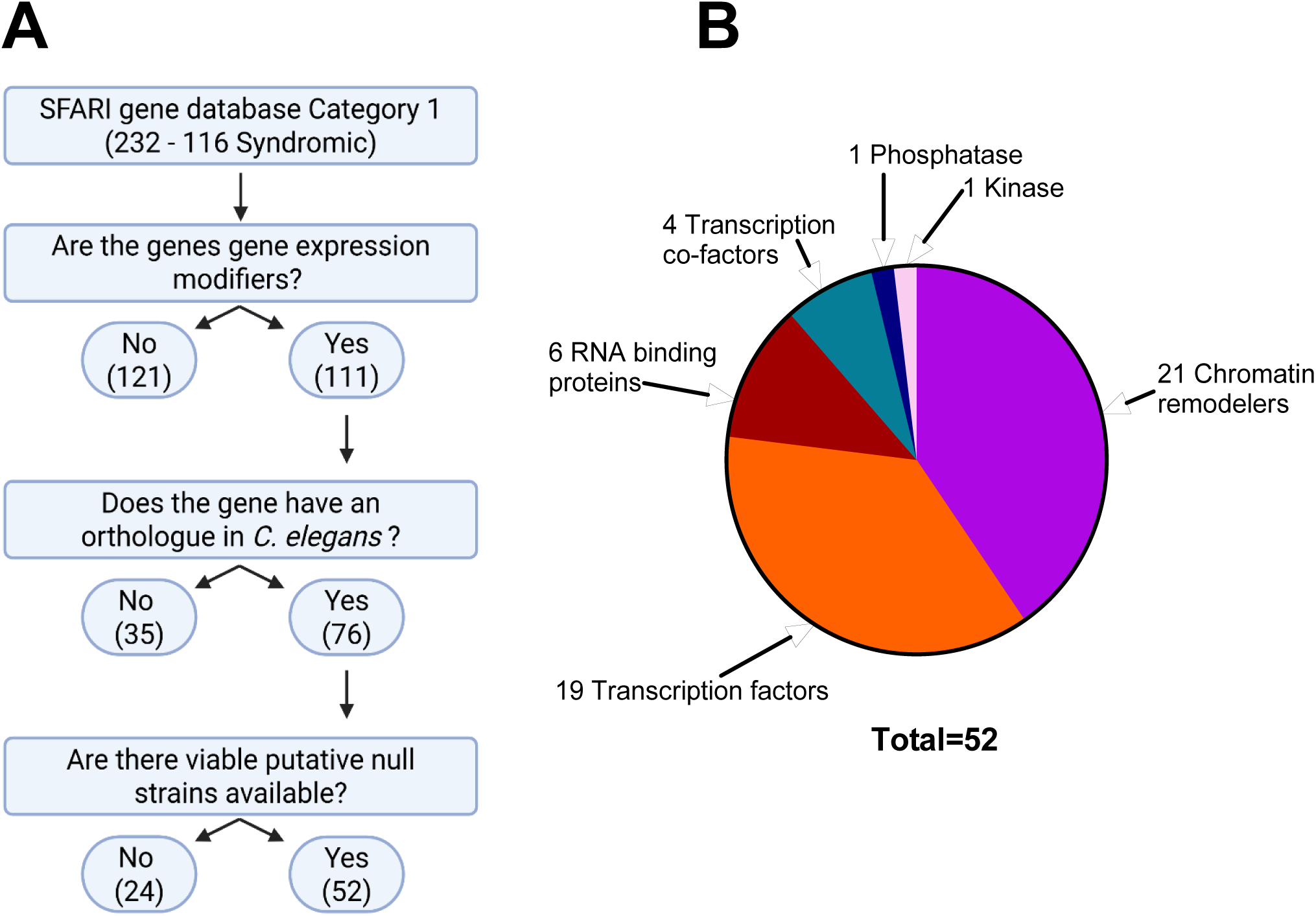
Schematic process and diagrammatic outcome of selected ASD-linked gene expression modifiers conserved in *C. elegans*. **A** Schematic diagram showing the bioinformatic filtering process of ASD related gene expression modifiers found in the SFARI gene database. The approach began with 232 total genes classified by SFARI as Category 1, of which 116 are further classified as syndromic. The number of genes selected at each point in the pipeline are indicated in brackets. Ortholist2 was used for determining orthologues of SFARI genes. Categories of worm genes were based upon the published role of the human orthologue. **B:** The breakdown of the 52 candidate genes based on their molecular function. Includes: chromatin remodelers, RNA binding proteins, transcription co-factors, phosphatase, kinase, and transcription factors. Molecular function was assigned based on the published role of their human orthologue or gene ontology (GO) with additional verification using available structural motif analysis.

*C. elegans* strains with putative null mutations in orthologous genes were identified (Table S2). Mutant alleles were documented as either large deletions or point mutations with a high variant effect predictor (VEP) score (Szklarczyk et al., 2023) indicating the probable loss of protein function by instability or nonsense mediated decay. We analysed each mutant on this basis without performing a mechanistic investigation into each gene. Mutant strains for 52 (68%) of the 76 orthologous genes identified were selected for further downstream analysis. Whilst mutant strains were available for the other 24 orthologous genes, they were not selected because the mutation generated either a lethal or sterile phenotype, making them unsuitable for our downstream analysis. In the case of some genes, up to two strains with distinct alleles were analysed. Additionally, of the 65 strains, 16 were outcrossed at least once but the remaining 49 were un-outcrossed and could limit interpretation of the data.

Each gene was assigned a molecular function identified from primary literature searches for the human orthologue. The molecular function of the 52 candidate genes can be classified into two primary groups; chromatin remodelers and transcription factors which make up 40% and 35% of the total respectively (Fig 1B).

RNA binding proteins and transcription co-factors contribute 13% and 8% of the total. Finally, a single phosphatase and kinase emerged through the selection pipeline described above. These were included in this analysis despite not directly driving transcription or translation functions due to their phosphorylating or dephosphorylating activity on gene expression modifiers (Atas-Ozcan et al., 2021; Smolen et al., 2023).

### Experimental pipeline for the phenotypic analysis of selected *C. elegans*

### mutants

A combination of developmental, anatomical and chemosensory driven behavioural assays was conducted on the selected *C. elegans* strains (Fig 2). These were based on robust assays which have already been validated for measuring either sensory or developmental phenotypes (Hilliard et al., 2002; Bargmann, 2006; Altun, 2009). Sensory assays were performed using the aversive cues: low pH, fructose, copper sulphate and octanol, whilst a low concentration of diacetyl was used as an attractive cue. These are established sensory cues known to drive behavioural responses in *C. elegans* that can be readily quantified.

**Fig 2.**
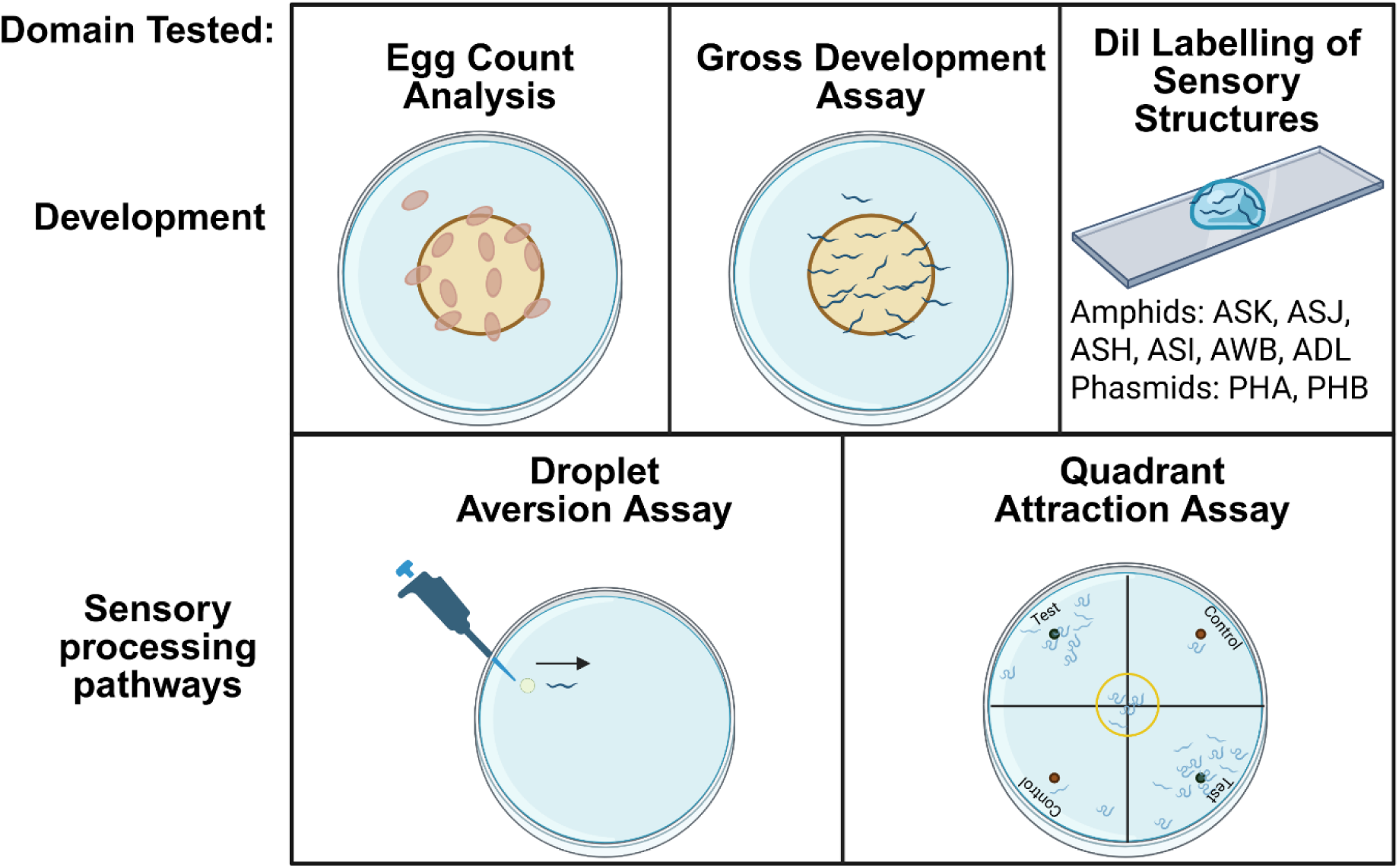
Assay schematics for interrogating the developmental and chemosensory capabilities of *C. elegans*. Assays used within this pipeline to interrogate the developmental and chemosensory phenotypes of *C. elegans* strains with ASD associated mutations. For a readout of reproduction an egg count analysis was used. For determining developmental capabilities, a gross developmental assay and DiI labelling of sensory amphids and phasmids were used. For determining sensory capabilities, a droplet aversion assay (aversive chemosensory) and quadrant attraction assay (attractive chemosensory) were used. Image produced using BioRender.

### ASD associated mutants show significant disparate effects on egg laying

28 mutants (43%) spanning all the functional subgroupings had a significantly lower egg laying rate than N2 (Fig 3). The lowest egg laying rate was observed for the mutant *egl-27 (ok1670)*, confirming previous observations (Trent et al., 1983). Only one mutant, *baz-2 (tm235)* encoding a chromatin remodeller had a significantly higher egg laying rate compared to the N2. Interestingly, chromatin remodellers were the least affected group with only 5 strains (19%) showing a significant difference compared to N2 (*xnp-1 (tm678), baz-2 (tm235), jmjd-3.1 (gk384), jmjd-3.1 (gk387), met-1 (tm1738)*). In contrast, 58% of transcription factors and 86% of RNA binding protein strains were significantly reduced in egg laying compared to N2. This assay can therefore identify phenotypes based on the integrative behaviour of egg laying from distinct molecular backgrounds and suggests any dysregulation of reproductive systems is more likely to have a negative impact on fecundity.

**Fig 3.**
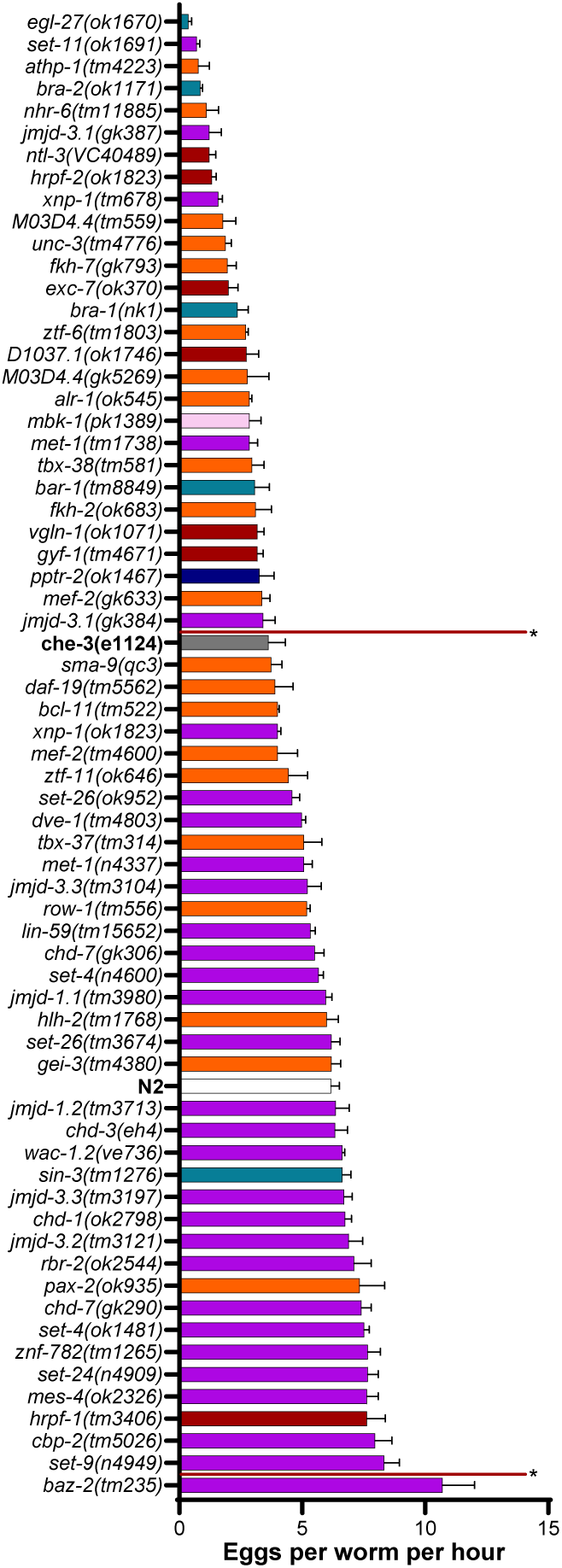
Egg count analysis of N2 vs ASD associated mutants. Egg count analysis comparing the egg laying rate of 3 worms over 3 hours between N2, ASD associated mutants and chemosensory deficient mutant *che-3 (e1124)*. All data is shown as mean ± SEM. Statistical analysis performed using one-way ANOVA with Dunnett’s multiple comparison test. n = 41 for N2 and n = 3 for all other mutants. Vertical red lines indicate significance compared to N2, strains not contained between the red lines are significantly different to N2. Chromatin remodelers, transcription factors, RNA binding proteins, transcription co-factors, phosphatase, kinase.

Whilst disruption to chemosensory processes has been previously described for mutants that have an impaired ability to detect food cues (L’Etoile et al., 2002; Chen et al., 2025), we did not identify a significant difference between the egg-laying rate of the *che-3 (e1124)* chemosensory deficient mutant and the wild-type. Furthermore, the egg-laying rate observed in some of the mutant lines was up to two-fold lower compared to the *che-3* mutant line and suggests the observed reduction is not fully explained by perturbation of sensory processes supported by *che-3* function.

### Pervasive and transient developmental delays observed in ASD associated mutants

Precise co-ordination of gene expression is important for tissue development and function. We measured the developmental progression of the selected ASD associated mutant strains by recording the proportion of developmental stages at timepoints in the *C. elegans* developmental lifecycle that corresponds to larval moults during early, mid and late development. At 20 hours, the approximate time of the L1 moult to L2, the proportion of L1 worms was 7% for the N2 as most of the population had undergone the first larval molt to L2 (Fig 4A). The chemosensory deficient mutant, *che-3 (e1124)* displays significantly delayed development across all life stages. Thus, chemosensory deficits can give rise to developmental delays which are pervasive providing a benchmark for solely chemosensory based developmental delays. The L1 proportion could be impacted by slow development during embryogenesis and/or during larval development. Unlike the egg count analysis, there was no clear association between a specific molecular function and phenotypic output (Fig 3, 4 A,B,C).

**Fig 4.**
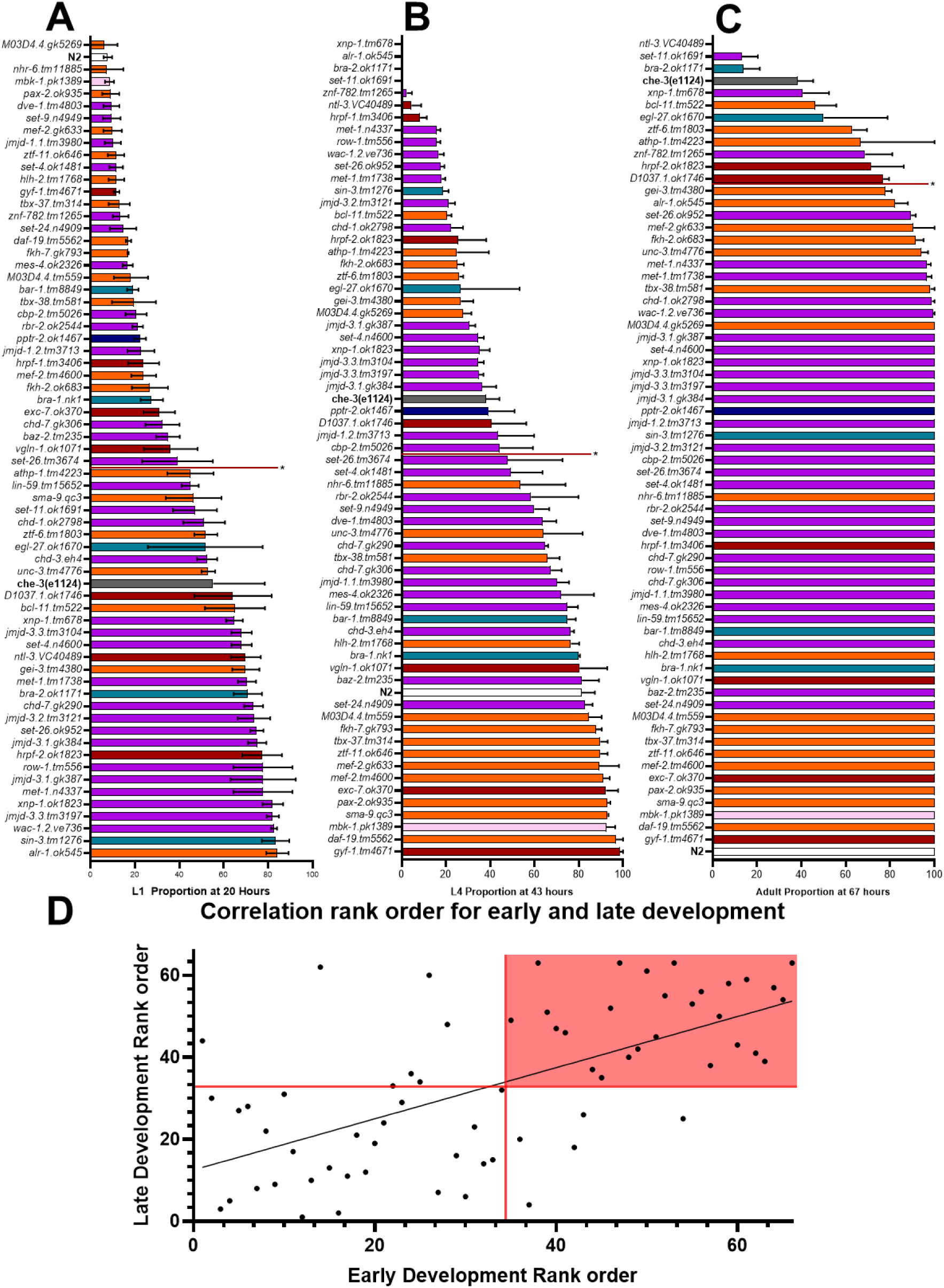
Gross developmental delays identified in ASD associated mutants vs N2. **A.** L1 proportion at 20 hours (approximate time of L1 moult to L2) of N2 compared to ASD associated mutants and chemosensory deficient mutant *che-3 (e1124)*. **B.** L4 proportion at 43 hours (approximate time of L3 moult to L4) of N2 compared to ASD associated mutants and *che-3 (e1124)* **C.** Adult proportion at 67 hours (sufficient time for N2 worms to reach adulthood) of N2 compared to ASD associated mutants and *che-3 (e1124)*. n = 18 for N2 and n = 3 for all other mutants. All data is shown as mean ± SEM. Statistical analysis performed using one-way ANOVA and Dunnett’s multiple comparison test. Vertical red lines indicate significance compared to N2, anything above the red line is significantly different compared to N2. Chromatin remodelers, transcription factors, RNA binding proteins, transcription co-factors, phosphatase, kinase. **D** Correlation of each strains developmental rank order identified by developmental rate analysis. Early development is defined as a significant difference from wild-type (N2) at L1/L2 moult occurring at ∼20 hours. Late development is defined as a significant difference from the wild-type (N2) at the L3/L4 moult time of ∼43 hours or delayed reaching adulthood ∼67 hours. The red lines mark the significance cutoff vs N2, and the red square highlights the area which is significantly different in both developmental categories.

Over half of the strains (34 of 65) investigated displayed an early developmental delay. A significant delay in early development was observed in the mutant strain *alr-1 (ok545)* (Fig 4A), which had the highest proportion of L1s compared to N2 (12-fold increase). A significant delay in mid and late development was observed in 33 of the 65 strains (Fig 4B, 4C). Plotting a rank order comparison of early and late development for each strain can show how pervasive the developmental delays are. There was a high concordance of strains with significant early and late developmental delays. This is indicated by 80% of strains with an early developmental delay also satisfying the criteria for delayed later stages of development (Fig 4D). This shows many strains fit the pattern observed in human cohorts of a pervasive developmental disorder which is present early in development and sustained (Minniscalco and Carlsson, 2022). However, 18% of the strains identified with a late developmental delay had no observed early developmental delay. One example of this is *M03D4.4 (gk5269)* (VEZF1 orthologue) which has a very similar proportion of L1s at 20 hours compared to N2 but a two-fold reduction in L4 proportion at 43 hours. Suggesting, that a developmental pattern is being perturbed between the L1 and L4 molt stages. Alternatively, 15% that had an early developmental delay showed amelioration by the time these strains reached late development. In this case it implies potential compensatory mechanisms within these strains. Finally, a large proportion encompassing 27 strains had no identified developmental delay.

### Changes in DiI labelled ciliated-sensory neuron number are rare in ASD associated mutants

In the context of ASD, the interaction between neurodevelopmental programming and structural changes in the nervous system are well documented (Hashem et al., 2020). To address if there was a detectable difference we focussed on a subset of sensory cells that mediate the response to environmental cues located in the amphid and phasmid structures. Given the growing emphasis of research on sensory systems in ASD, we wanted to develop the analysis around assays that quantified responses to distinct chemosensory cues. These environmental inputs are mediated primarily by amphid and phasmid neurons. When dysfunctional, they underpin disrupted sensory responses. Before initiating assays to quantify chemosensory responses, we first investigated the structural integrity of a subset of sensory amphid neurons and both phasmid neurons using DiI staining (as described in Figure 2).

The DiI staining was used to test for the presence of neurons primarily associated with sensory detection that lie upstream of the integrating and output components of the circuit that co-ordinate integrated behaviour. As previously described, DiI staining was sufficient to detect the cell bodies and the gross morphology of the 6 amphid sensory neurons ASK, ASJ, ASH, ASI, AWB and ADL and both phasmid neurons, PHA and PHB in the N2 strain (Fig 5 A, B) (Collet et al., 1998). The same staining protocol was applied to mutants and then imaged for amphid and phasmid cell bodies and gross morphology. The mutant line *che-3 (e1124)*, which is deficient for DiI staining (Wicks et al., 2000) was included in the analysis as a positive control (Fig 5 C, D).

**Fig 5.**
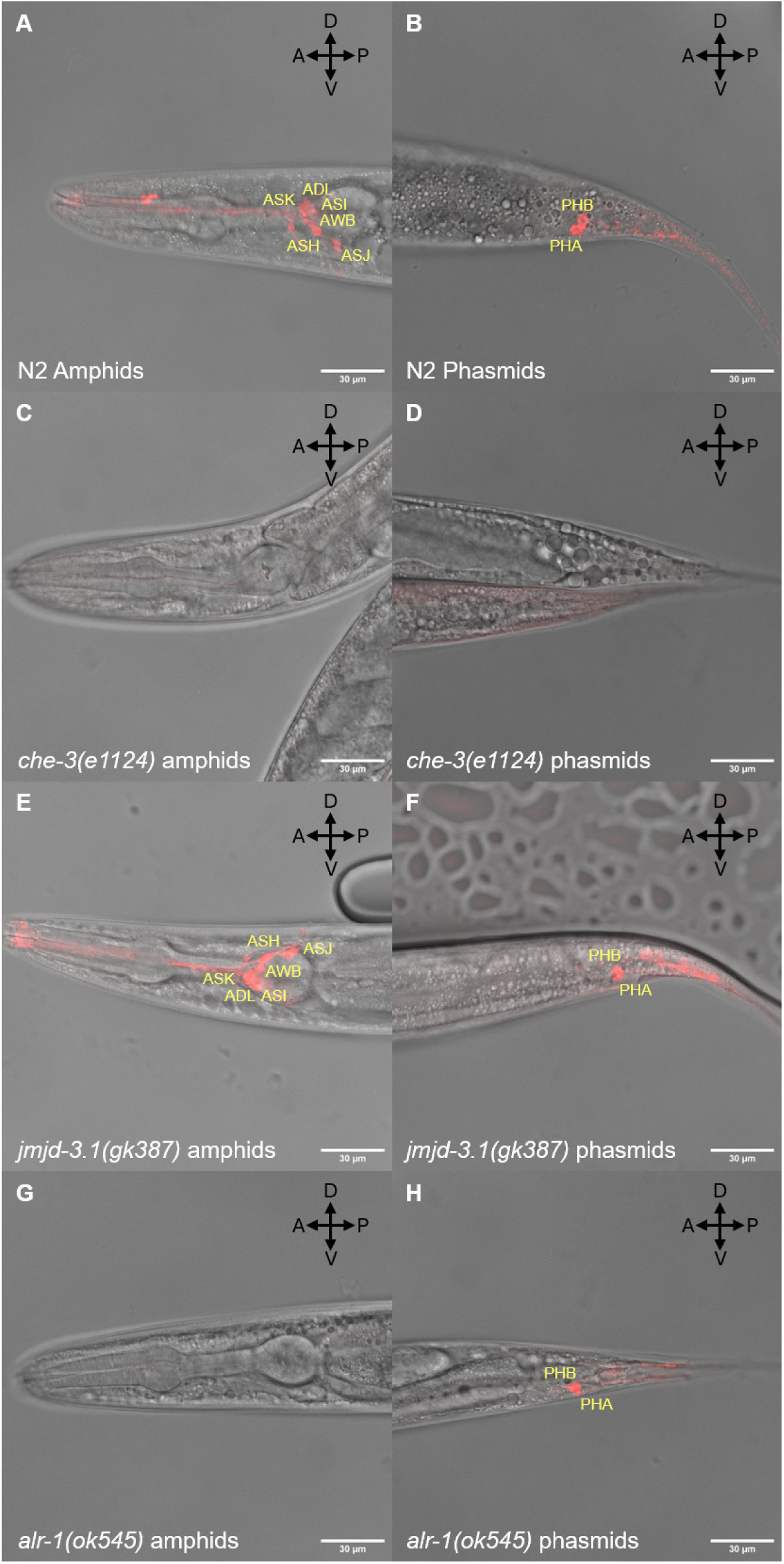
DiI staining of sensory neurons in ASD associated *C. elegans* mutants. **A.** Composite image of N2 head showing sensory amphids overlaid with DIC. Image labelled with visible amphid neurons. **B.** Composite image of N2 tail showing sensory phasmids overlaid with DIC. Image labelled with visible phasmid neurons. **C.** Composite image of *che-3 (e1124)* head showing missing sensory amphids overlaid with DIC. **D.** Composite image of *che-3 (e1124)* tail, overlaid with DIC showing an absence of DiI labelling of phasmid neurons **E.** Composite image of *jmjd-3.1(gk387)* head showing sensory amphid neurons overlaid with DIC. Image labelled to indicate visible amphid neurons. **F.** Composite image of *jmjd-3.1 (gk387)* tail showing sensory phasmid neurons, overlaid with DIC. Individual, visible phasmid neurons are labelled. **G.** Composite image of *alr-1 (ok545)* head overlaid with DIC. **H.** Composite image of *alr-1 (ok545)* tail showing sensory phasmid neurons overlaid with DIC. Individual, visible phasmid neurons are labelled. All images oriented as left – anterior, right – posterior, top – dorsal and bottom – ventral. DiI labelled structures are shown in red.

Only two of the ASD associated mutants had an alteration to the number of amphid/phasmid neuron cell bodies and gross morphology detected by the DiI, consistent with previously reports. These mutants were *alr-1 (ok545)* (Tucker et al., 2005) and *egl-27 (ok1670)* (Solari et al., 1999). For *alr-1 (ok545)*, none of the six sensory amphid neurons were detected by the DiI in the adult worm (Fig 5G).

However, this was selective as phasmid neurons were still detected by dye filling (Fig 5H) in two of the six worms imaged. In the case of *egl-27 (ok1670)*, none of the phasmid neurons were detected by the DiI but all six sensory amphid neurons could be visualised with the DiI. These observations are consistent with previous descriptions of DiI staining performed on *egl-27* and *alr-1* mutant strains (Tucker et al., 2005; Melkman and Sengupta., 2005; Trent et al., 1983; Desai et al., 1988).

Thus, the presence of sensory neurons was unaffected in most of the strains when compared to N2. For example, the cell bodies of all amphid and phasmid neurons stained by DiI could be detected in the strain *jmjd-3.1 (gk387)* (Fig 5 E, F) but as we describe later, this strain exhibited both pervasive developmental delays as well as sensory deficits. Hence, the mutants in the pipeline showed little impact on the complete loss of neurons that initiate sensory processing, thus informing interpretation of subsequent behavioural analysis. Our analysis was limited to counting neuron number and assessing gross neuron morphology. The possibility of more subtle morphological changes in the cilium or sensory neuron akin to those previously implicated in ASD (Martínez-Cerdeño et al., 2016) could account for the phenotypic observations and cannot be excluded.

### ASD associated mutants cause hypo or hypersensitive behavioural responses to chemosensory stimuli

We evaluated chemosensory processing capabilities within each strain using chemoattractant and aversive cues. A diacetyl (1% v/v) quadrant assay (Fig 2) was utilised to measure attractive chemosensory processing. In this assay each worm had to detect the positive cue and integrate that signal into a locomotory response to move in the direction of the cue (chemotaxis). The observed chemotaxis of the mutants was compared to the N2 performance to determine any modified sensitivity to the attractive cue.

In addition, aversive chemosensory processing was assayed by quantifying reversals to both water soluble and volatile aversive cues. Processing of attractive and aversive chemosensory cues requires the functionality of different sensory amphid neurons (Ferkey et al., 2021), but downstream interneuron and motor neuron circuits are shared. Thus, by comparing strains across multiple chemosensory cues it is possible to distinguish between strains impacted at the level of sensory detection versus those impacted in downstream circuitry.

#### Chemotaxis response perturbed in 25% of ASD associated mutants

N2 showed a reproducible attraction to the chemosensory cue diacetyl (1% v/v) as evidenced by the performance index of 0.8 (Fig 6A). 17 mutants showed a significant reduction in their performance index compared to N2. There was no clear association with a single molecular functional group, with each group represented proportionally. Additionally, only one strain, *M03D4.4 (tm559)*, showed a performance index lower than the negative control *che-3 (e1124)* of 0.05 with the remaining significantly impaired strains scoring between 0.07 and 0.42. This suggests an incomplete loss of sensory detection or sensory processing as most mutants fall between normal processing by N2 and complete loss of sensory processing by *che-3 (e1124)*.

**Fig 6.**
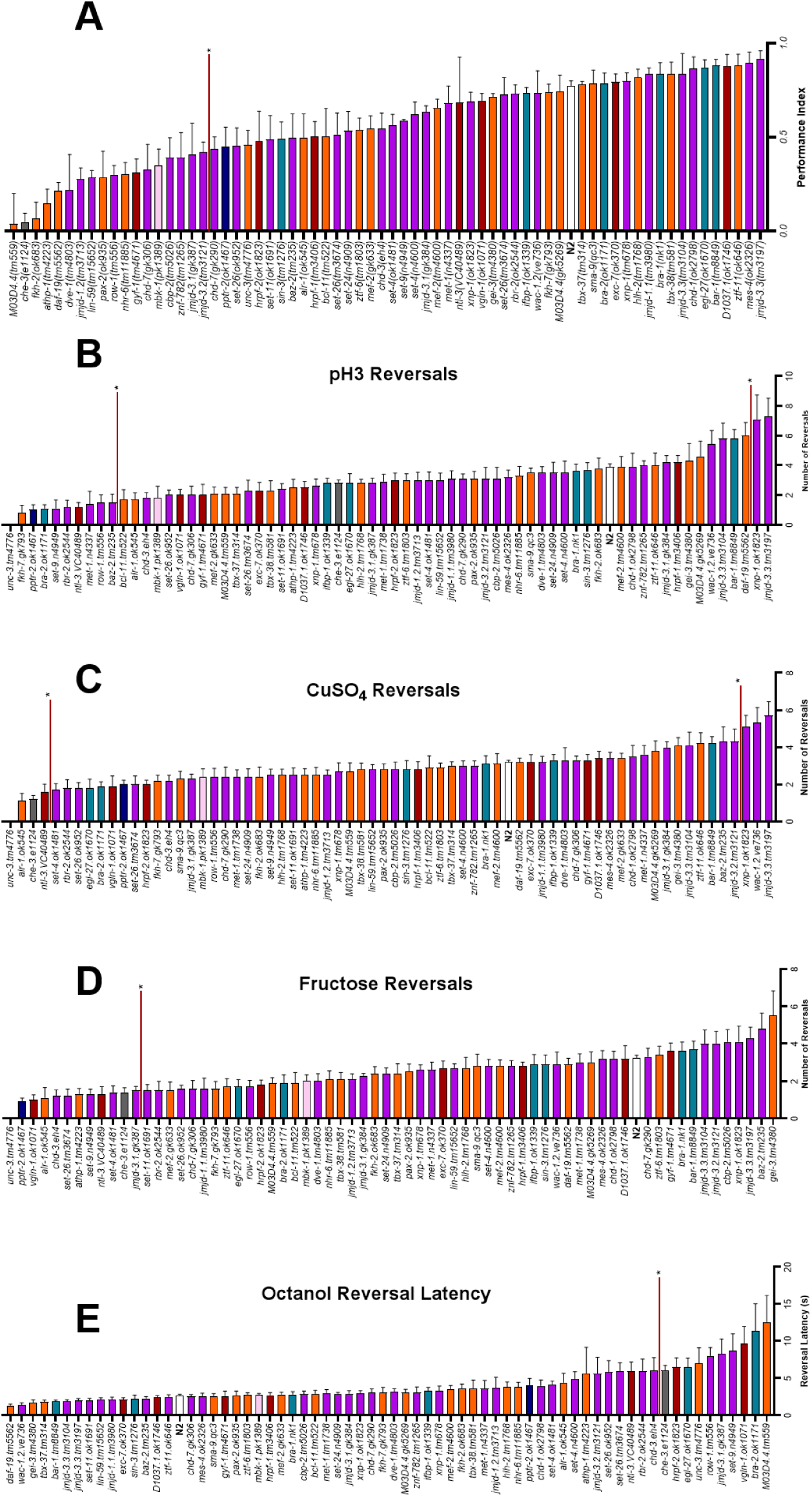
Sensory processing capabilities modified in *C*. *elegans* ASD associated mutants. **A.** Performance index of N2 vs ASD associated mutants in diacetyl quadrant assay. n=48 for N2. n = 3 for all other strains. Statistical analysis performed using one-way ANOVA and Dunnett’s multiple comparison test. **B, C, D.** Average number of reversals of N2 vs ASD associated mutants in response to aversive stimuli (low pH, copper sulphate and fructose). **E.** Latency to initiating reversals of N2 vs ASD associated mutants in response to 30% octanol. **B, C, D, E.** Statistical analysis performed using one-way ANOVA and Dunnett’s multiple comparison test. n = 180 for N2 and n = 10 for all other mutants. All data is shown as mean ± SEM. Chromatin remodelers, transcription factors, RNA binding proteins, transcription co-factors, phosphatase, kinase. Significance shown by red horizontal line, in A and D strains to the left of the red line are significant in B and C the strains outside of those contained between the red lines are significant vs N2 and in E, the strains to the right of the red line are significantly different to N2.

#### Aversive cue responses show hypo and hypersensitivity

We measured responses to different aversive cues which utilise distinct molecular determinants to trigger the reversal behaviour. In addition, we extended this to aversive volatile cues. Hypersensitivity, defined in this study as an enhanced aversive behavioural response compared to the wild-type was observed in *xnp-1* (*ok1823*), *jmjd-3.3* (*tm3197*) and *wac-1.2* (*ve736*) in response to low pH and copper sulphate (Fig 6B, 7C) and 1-octanol (Fig 6E). Hyposensitivity, defined as a reduced aversive behavioural response was exhibited by 17 mutants, to one or more of the soluble aversive cues. 6 of the 9 strains that were hyposensitive to 1-octanol also demonstrated hyposensitivity to at least one other soluble cue (Fig 6B-E) (*jmjd-3.1 (gk387)*, *set-9 (n4949), row-1 (tm556), unc-3 (tm4776), vgln-1 (ok1071), bra-2 (ok1171)*).

Of the 65 strains tested, 35 reached the criteria that indicate they show significant changes in response to the sensory cues (Fig 6B-E) compared to N2.

Hyposensitivity to the aversive cues was more commonly observed at 30 instances compared to just 5 instances of hypersensitivity.

### The significance heatmap of developmental and sensory assays of ASD associated mutants reveals functional segregation

The results of the developmental, anatomical and chemosensory phenotypic analysis are summarised in figure 7 and 8. Each row is the specific phenotype observed in a particular strain and can be directly compared to other strains to identify phenotypic overlap within the dataset (alternative grouping in Fig S2). Furthermore, each gene is associated with a specific molecular functional grouping allowing for comparisons across individuals within a group as well as across groups.

**Fig 7.**
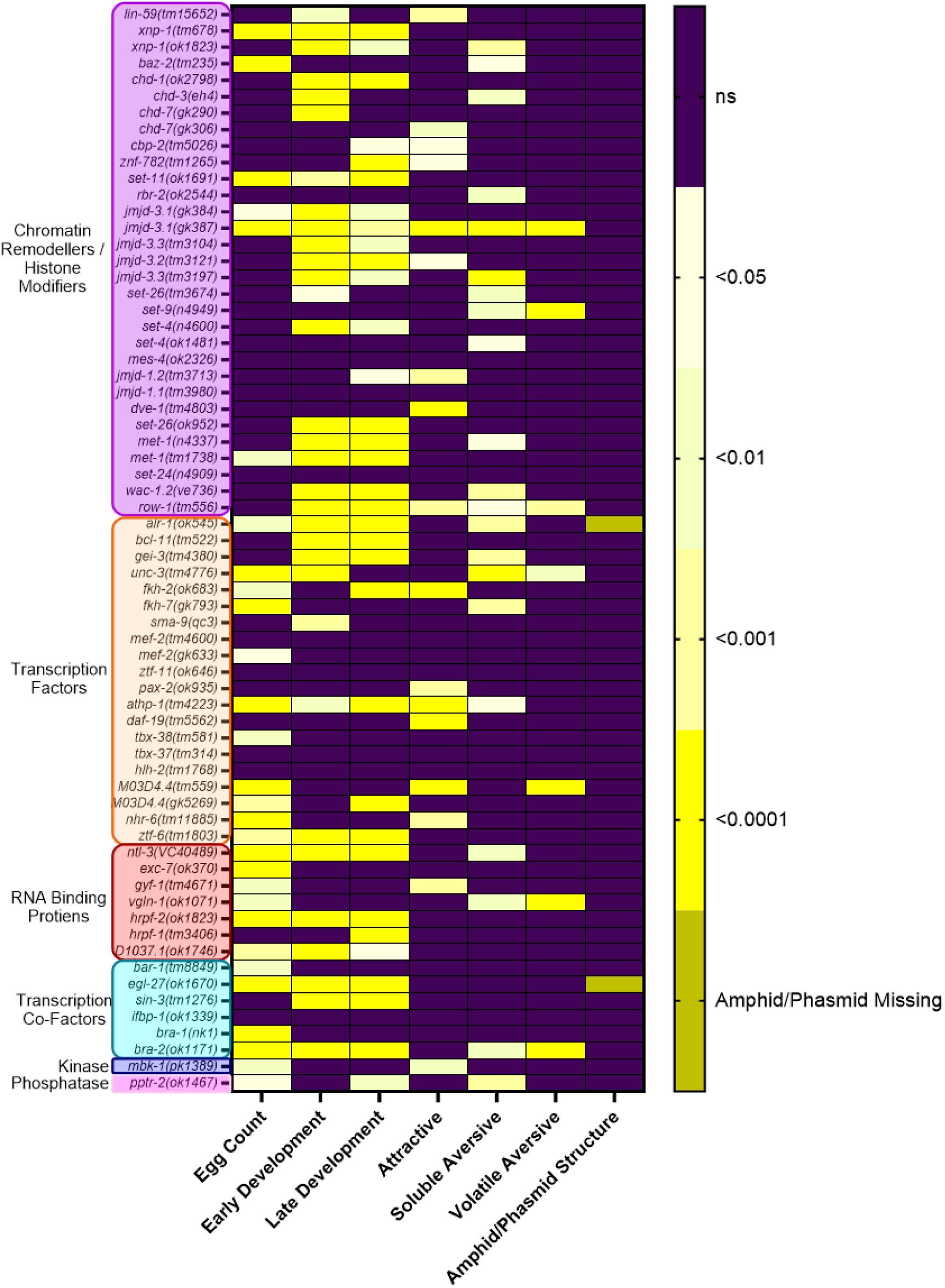
Significance heatmap of ASD associated mutant’s vs N2 for developmental and sensory phenotypes. Significance heatmap showing the seven phenotypes assessed in the experimental pipeline including, egg count, early and late development, attractive chemosensory processing, soluble aversive chemosensory processing, volatile aversive chemosensory processing and sensory amphid/phasmid structure. Yellow boxes show significant difference from N2, and purple are non-significant compared to N2, in each respective phenotype. Significance is defined at p≤0.05. For egg count, early and late development, attractive, soluble aversive and volatile aversive chemosensory processing one-way ANOVA and Dunnett’s multiple comparison test was used for statistical analysis. For amphid/phasmid DiI staining any deviation from the expected six amphids and two phasmids was noted as significant.

Egg laying appears to be very sensitive to mutant perturbation, with over 50% of transcription factors and over 80% of RNA binding proteins showing significance compared to less than 20% of chromatin remodellers studied (Fig 7). Early and late developmental phenotypes are not associated with any one molecular functional grouping as evidenced by a consistent percentage of strains in each group showing developmental delays (Fig 7). Furthermore, there are no genes that are significantly affected across all phenotypes suggesting that there is not a fully penetrant sensory deficient mutant that is also impaired in development and DiI labelling. Concordance between the 9 pairs of alleles was 70% across all phenotypes with over 50% of discrepant phenotypes being sensory processing.

Across all strains analysed 89% had at least one phenotype (Fig S1A) leaving just 7 strains (*mes-4 (ok2326), jmjd-1.1 (tm3980), set-24 (n4909), mef-2 (tm4600), tbx-37 (tm314), hlh-2 (tm1768, ifbp-1 (ok1339)*) which had no phenotype across any of the assays used here. Gross changes to sensory structures were the least commonly identified phenotype affecting just 2 strains (*alr-1* and *egl-27)* both of which showed additional phenotypes in development and sensory processing (Fig 8).

**Fig 8.**
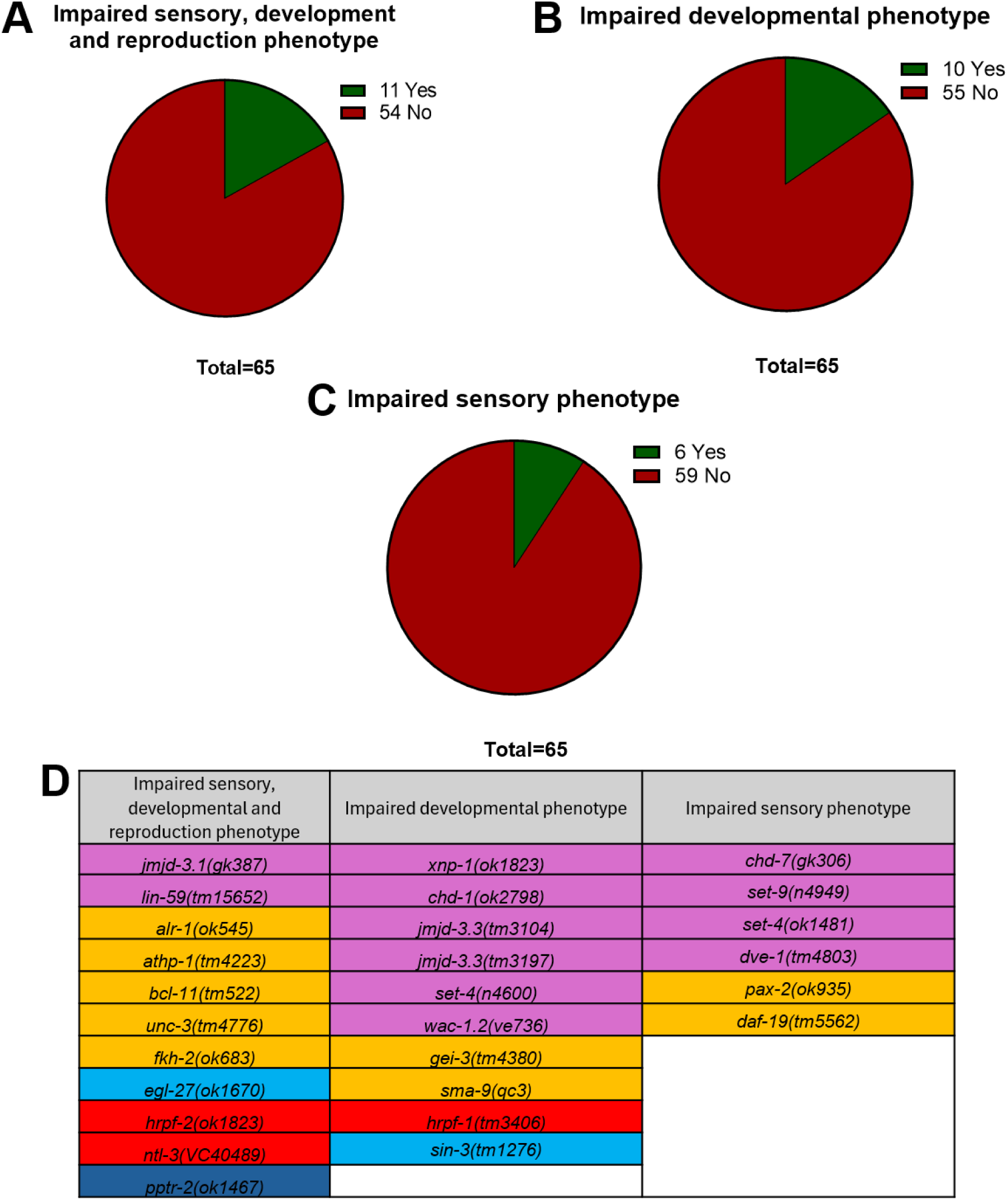
Filtering for significant developmental or sensory affected mutants highlights three distinct groupings. **A.** Subclassification of mutant strains for deficits in sensory processing, gross developmental delays and egg laying deficits. **B.** Phenotypically grouped genes for deficits in developmental programs but without any sensory function deficits. **C.** Phenotypically grouped genes for deficits in sensory function without any gross developmental delays or reproductive deficits. **D.** Genes from the significance heatmap grouped according to either impaired sensory, impaired developmental or a combination of phenotypes. The phenotypes grouped as developmental for the purpose of filtering were egg count, early development and late development. The phenotypes grouped as sensory were attractive chemosensory, soluble and volatile aversive chemosensory.

Developmental or sensory phenotypes were almost equally common in the strains investigated at 42 and 38 strains respectively (Fig S1A). Finally, chemosensory impairment overlapped with both developmentally delayed and non-delayed strains at a similar percentage but there is a prominent difference in the overlap of severe chemosensory impairment with developmental delay compared to non-delayed strains (Fig S1B).

### Three phenotypic groupings emerge from developmental and chemosensory screening of ASD associated mutants

We conducted a phenotypic grouping analysis to parse the cohort of mutants that passed through the comparative pipeline. This approach, that integrates phenotypic score of each gene, established 3 groups of genes. The classification was driven by ascribing normal or perturbed phenotypic response based on multiple comparison statistical tests of each gene in each assay.

The subgroups derived from Figure 8 were classified as having either a sensory phenotype or a developmental phenotype or a combined sensory, developmental and reproductive phenotype relative to the wild-type control. Group 1, deficient in both sensory responses, development and reproduction, includes 2 chromatin remodelers: 5 transcription factors and one cofactor, 2 RNA binding protein and a phosphatase (Fig 8 A, D). Group 2, impaired in only development, includes 6 chromatin remodelers, 2 transcription factors, one cofactor and an RNA binding protein (Fig 8 B, D). Group 3, impaired in only sensory responses, includes 4 chromatin remodelers and 2 transcription factors (Fig 8 C, D). Group 1 genes show a significant difference across all phenotypes investigated except changes in DiI labelling which provides a subset of genes with pleiotropic effects. Group 2 and 3 genes show more selective effects in development and chemosensory processing respectively.

## Discussion

A widely recognised phenotypic trait associated with ASD is atypical sensory processing that can be readily observed in clinical cohorts and has emerged as a listed characteristic in the updated DSM V (American Psychiatric Association, 2013). This supplements the established criteria around social and repetitive motor behaviours and the characteristic early life emergence of the neuroatypical traits. In clinical cohorts the reported changes span multiple sensory modalities including visual, auditory, tactile and chemosensory (Sivapalan et al., 2024; Bennetto et al., 2007), although they are not systematically recorded (Table S1). However, individuals with ASD have shown an increased severity in behavioural challenges because of sensory processing changes requiring routine therapy. (Sivapalan et al., 2024). This highlights the severe challenges for individuals with ASD and the lack of genetic or pharmacological therapeutic targets available.

We wanted to investigate how sensory functions are perturbed by designing a pipeline in *C*. *elegans* to exploit approaches that inform on this aspect. We designed the pipeline to resolve epigenetic and wider genetic modulators in which there is good evidence for intersection with environmental cues that go through sensory pathways (Peña., 2025; Pumo et al., 2022). In addition, this pipeline utilises epigenetic regulators that contribute to polygenic transcriptional programs that may represent a distributed polygenic landscape like those that represent the most significant genetic determinants of ASD. The pipeline is supplemented by additional transcriptional regulators. In restricting our analysis to category 1 SFARI genes, of which half are further classified as syndromic, it allowed the comparison of key biological traits encompassing developmental and sensory responses in the highest confidence gene expression modifiers.

Furthermore, developmental delays in ASD cohorts have been correlated with a greater likelihood of other ASD symptoms such as impairments in fine motor skills, language ability and personal social activities (Shan et al., 2022). For this analysis, it is important to note that intrinsic changes in gross development represented in the mutations are confounded by the potential for gross sensory disruption which is sufficiently penetrant to drive developmental changes as exemplified by the positive control *che-3 (e1124).* Therefore, understanding the intersection of developmental processes and sensory processing changes in ASD is vital to develop pharmacological treatments.

We identified that development and by extension underpinning programs were especially susceptible to perturbation, with 61% of strains being impacted in either early or late development. Subtle transitions occur between early and late larval stages and by the fourth molt, 400 epidermal, gonad, muscle and neuronal cells have been added to a total of 959 somatic cells (Rougvie and Moss, 2013). This process is susceptible to perturbation at any point which could lead to a developmental delay.

Mutants that showed perturbations early in development were largely pervasive with over 80% concordance of strains with significant early/late developmental delays suggesting delays in early development were not easily overcome. However, this was not universal and 15% of strains with an early developmental delay showed no significant delay in later development, suggesting a possible mechanism by which these strains were able to expedite their later development. This is not unprecedented as early perturbation of developmental programs have been corrected by late expression of the affected gene (Luikenhuis et al., 2004).

One interesting possibility for comparison to the human disorder can be made here because spontaneous recovery in autism associated phenotype has been reported and suggests the possibility that recovery of developmental delays is co-ordinated through more widely shared biological mechanisms (Sitholey et al., 2009). Another example is MeCP2 the main driver of Rett’s Syndrome (RTT) where restoring the expression of a functional MeCP2 rescues the RTT phenotype in a mouse model (Luikenhuis et al., 2004). Despite this phenotypic rescue originating from transgenic expression of the affected gene, it does not preclude the possibility for endogenous mechanisms being present within *C. elegans* that are capable of self-rescuing developmental delays. Additionally, environmental cues from other worms on the same plate have been shown to affect developmental rates, providing an example of environmental impact on development which could also contribute to the developmental phenotypes we observed (Ludewig et al., 2017).

The criteria used to define chemosensory dysfunction in our study was a significant difference compared to N2 in either attractive, soluble aversive or volatile aversive chemosensory processing. However, when considering mutants that had impaired development, the potential impact becomes clearer when highlighting a more severe sensory phenotype. Increased severity here is defined as significant perturbations across multiple chemosensory cues. The severity of sensory perturbation overlaps with developmental delays, supporting our approach of scoring development alongside behavioural outcomes because 28% of strains with an impairment in early or late development also show a more severe impact on sensory processing.

Interestingly, it is only the severity of the sensory perturbations which correlates with developmental delays, as developmentally delayed strains had a 55% overlap with sensory perturbations compared to 60% of non-developmentally delayed strains.

This highlights the worm has a distinct relevance to modelling this condition as the same effect of developmental delays correlates with greater severity of ASD symptoms in human cohorts (Shan et al., 2022). Furthermore, it shows that sensory perturbations in the adult can arise without a developmental delay being present and suggests the possibility for selectivity in the genetic underpinnings of different subsets of ASD mutants. This provides further evidence for severe sensory deficits leading to developmental delays (Fujiwara et al., 2002; Rose et al., 2005) although the direct cause of developmental delay is not investigated here.

Attractive and aversive chemosensory cues can be detected by distinct amphid neurons coupled to a common downstream signalling circuitry, comprised of shared interneurons (White et al., 1986; Larsch et al., 2015; Piggott et al., 2011). Hence, disruption to the signalling circuitry downstream of the amphid neurons would be expected to disrupt responses to both attractant and aversive chemosensory cues. In our analysis, disrupted attractive and aversive chemosensory phenotypes combined in 3 out of the 35 total strains that exhibited a chemosensory deficit (*athp-1 (tm4223), row-1 (tm556)* and *jmjd-3.1 (gk387)*). This observation is consistent with a disruption to shared components of the circuitry downstream of the amphid neurons. Further, as most of the sensory deficits observed are either to aversive or attractive cues it supports that in most cases, the chemosensory impairment observed is likely to be at the level of detection and localised to the amphid neurons. Selective differences could be caused by differential expression of G protein-coupled receptors (GPCRs) or associated signalling apparatus which has been implicated in ASD aetiology (Hormozdiari et al., 2015; Monfared et al., 2021).

Our analysis led to three groups of mutants which can be aligned with dysfunction in different areas of ASD symptomology, namely Group 1: sensory, gross development and reproduction; Group 2: gross development and reproduction; Group 3: sensory. RNAi and genetic screens have been widely used in *C. elegans* to identify genotype/phenotype interactions allowing for comparisons of the developmental and where possible, reproduction and sensory phenotypes investigated here. For example, of the 11 genes in group 1, 4 have previously been shown to have slow growth and egg laying deficits (Ceron et al., 2007; Schmitz, Kinge and Hutter., 2007; Simmer et al., 2003; Shepard et al., 2011), 4 with no slow growth (Gönczy et al., 2000; Kamath et al., 2003) and only one has been previously described as having sensory deficits (Kim et al., 2005). By comparison, in group three (mixed phenotype) genes only 1 has previously been ascribed a slow growth phenotype (Kamath et al., 2003) but 3 have known sensory phenotypes (Mcdiarmid et al., 2019; Sheng et al., 2024; Swoboda, Adler and Thomas., 2000). For the three groups of genes identified here, overlap with reported phenotypes was 58% for group 1, 17% for group 2 and 90% for group 3. The amount of overlap with previously reported phenotypes could reflect the high rate of false negatives associated with RNAi screens, which is reportedly 59% (Rual et al., 2004).

The three sub-groups which are criteria-based, contain orthologues spanning diverse molecular functions. We speculate that these represent distinct but definable groupings in which the identified molecular perturbation generates shared mechanistic behaviours. This view expresses outcomes of the genetic interrogation of symptomology from human cohorts (Litman et al., 2025; Leyhausen et al., 2025).

These studies highlight the preferred view that distinct subcategories relate to distributed mechanistic underpinnings (Litman et al., 2025; Leyhausen et al., 2025). Our criteria-based approach in which the intrinsic developmental programs and associated behavioural outcomes in the adult are similar to a biologically simpler organism demonstrates the possibility of modelling specific ASD phenotypes like sensory dysfunction in *C. elegans*. This provides a basis for further analysis to investigate the transcriptional profile within each grouping identified in our study to determine if shared mechanistic pathways underpin behavioural changes within each group and differentiate the groups from each other.

These notions are consistent with recent evidence for selective pathways altered in ASD individuals creating a unique presentation of the disorder (Litman et al., 2025). However, one understandable limitation in these human cohort studies is the absence of sensory phenotypes in any of the groups they identify. The preferred interpretation of our data supports the need for a more systematic sensory testing of ASD cohorts. This would require quantitative sensory testing in ASD cohorts with known genetic backgrounds. Using the 3 gene groups segregated from our investigation we attempted to identify sensory phenotypes in human and mouse cohorts but due to limited sensory information it was difficult to determine if our identified genes correlated with human or murine findings (Table S1).

Our approach highlights that quantitative phenotyping can frame subcategorisation but is currently limited in defining underlying mechanism. An important task will be to develop this work to probe if discrete structural or molecular changes can be identified within distinct *C. elegans* behavioural subclassification. If these sub-classifications organised around investigating autism related genes, associate with discrete mechanistic underpinnings, it would encourage the view that the complexity of human neuroatypical spectrum disorders can be organised into dysfunctional sub-types to the benefit of diagnosis and mitigation.

In conclusion, investigation of ASD associated gene expression modifier orthologues in *C. elegans* led to a criteria-based grouping with shared phenotypes. These groupings encompass genes with distinct and divergent biological functions with a shared ability to modulate wider gene expression. We drove this by considering the phenotypic overlap between developmental timing and quantifiable, systematic measurement of sensory responses. The latter is an important and underreported component of ASD pathology with a direct impact upon an individual’s behaviour and quality of life (Giacardy et al., 2018). Therefore, further investigation into the molecular basis of sensory processing perturbation is necessary and our robust experimental pipeline allows the identification of candidate genes for further analysis.

