## Supplemental Figures for "Sensory and developmental phenotyping of *C. elegans* parses autism associated genes into behavioural classifications"

| Phenotypic Group | Strain | Human Gene | Syndromic? | Sensory phenotype in human cohorts? | Sensory phenotype in mouse models? |
|---|---|---|---|---|---|
| Impaired Sensory | *chd-7(gk306)* | CHD8 | YES | Not Reported | YES |
| | *set-9(n4949)* | KMT2E, SETD5 | YES x2 | Not Reported | YES |
| | *set-4(ok1481)* | KMT5B | NO | Not Reported | Not Reported |
| | *dve-1(tm4803)* | SATB1 | YES | YES | N/A |
| | *pax-2(ok935)* | PAX5 | NO | YES | Not Reported |
| | *daf-19(tm5562)* | RFX3 | NO | YES | Not Reported |
| Impaired development | *xnp-1(ok1823)* | ATRX | NO | YES | YES |
| | *chd-1(ok2798)* | CHD2 | YES | Not Reported | Not Reported |
| | *jmjd-3.3(tm3104)* | KDM6B | NO | Not Reported | Not Reported |
| | *jmjd-3.3(tm3197)* | KDM6B | NO | Not Reported | Not Reported |
| | *set-4(n4600)* | KMT5B | NO | Not Reported | Not Reported |
| | *wac-1.2(ve736)* | WAC | YES | YES | Not Reported |
| | *gei-3(tm4380)* | CIC | NO | Not Reported | Not Reported |
| | *sma-9(qc3)* | HIVEP2 | YES | Not Reported | Not Reported |
| | *sin-3(tm1276)* | SIN3A | YES | YES | Not Reported |
| | *hrpf-1(tm3406)* | HNRNPH2 | NO | Not Reported | Not Reported |
| Mixed | *jmjd-3.1(gk387)* | KDM6B | NO | Not Reported | Not Reported |
| | *lin-59(tm15652)* | ASH1L | NO | YES | Not Reported |
| | *alr-1(ok545)* | ARX | YES | YES | YES |
| | *athp-1(tm4223)* | PHF12 | NO | Not Reported | Not Reported |
| | *bcl-11(tm522)* | BCL11A | YES | Not Reported | Not Reported |
| | *unc-3(tm4776)* | EBF3 | YES | YES | Not Reported |
| | *fkh-2(ok683)* | FOXG1 | YES | Not Reported | YES |
| | *egl-27(ok1670)* | RERE | YES | YES | Not Reported |
| | *ntl-3(VC40489)* | CNOT3 | YES | YES | YES |
| | *hrpf-2(ok1823)* | HNRNPH2 | NO | Not Reported | Not Reported |
| | *pptr-2(ok1467)* | PPP2R5D | YES | YES | Not Reported |

**Table S1. Clinical sensory data of ASD associated genes matched to *C. elegans* phenotypic groupings**

ASD associated genes grouped phenotypically from an experimental pipeline of reproduction, development and sensory phenotypes. Human orthologues are displayed and matched to criteria of being syndromic, human sensory phenotypes and mouse sensory phenotypes. Phenotypic information gained through phenotypic databases, Human Phenotype Ontology (https://hpo.jax.org/), International Mouse Phenotyping Consortium (https://www.mousephenotype.org/), PUBMED searches for “Gene Name” AND “Sensory” AND “Human” OR “Mouse”. Sensory phenotypes were then recorded as yes or not reported if no evidence for sensory phenotyping exists.

| Human gene | Gene name | Protein function | C. elegans orthologue | Allele | Strain name | Outcrossed | Mutation | Egg Laying Phenotype | Slow Growth (Developmental) Phenotype | Sensory Phenotype | Dye Filling Phenotype |
|---|---|---|---|---|---|---|---|---|---|---|---|
| ASH1L | ASH1 like histone lysine methyltransferase | Histone methyltransferase | lin-59 | tm15652 | FX25652 | x0 | Deletion | Yes (Schmitz, Kinge and Hutter., 2007) | Yes (Schmitz, Kinge and Hutter., 2007) | Not Reported | Not Reported |
| ATRX | ATRX chromatin remodeler | SWI/SNF chromatin remodeler | xnp-1 | tm678 | FX00678 | x0 | Deletion | Not Reported | No ( Eki et al., 2007) | Not Reported | Not Reported |
| | | | | ok1823 | VC2088 | x1 | Deletion | | | | |
| BAZ2B | Bromodomain adjacent to zinc finger domain 2B | Component of chromatin remodeling complexes | baz-2 | tm235 | FX00235 | x0 | Deletion | Not Reported | No (Kamath et al., 2003) | Not Reported | Not Reported |
| CHD2 | Chromodomain helicase DNA binding protein 2 | SNF2 related chromatin remodeler | chd-1 | ok2798 | RB2613 | x0 | Deletion | Not Reported | No (Fraser et al., 2000) | Not Reported | Not Reported |
| CHD3 | Chromodomain helicase DNA binding protein 3 | SNF2 related chromatin remodeler | chd-3 | eh4 | HX103 | x4 | Deletion | Not Reported | No (Kamath et al., 2003) | Not Reported | Not Reported |
| CHD8 | Chromodomain helicase DNA binding protein 8 | SNF2 related chromatin remodeler | chd-7 | gk290 | VC606 | x0 | Deletion | Not Reported | No (Eki et al., 2007) | Yes (Mcdiarmid et al., 2019) | Not Reported |
| | | | | gk306 | VC676 | x0 | Deletion | | | | |
| CREBBP | CREB binding protein | Histone acetyltransferace and transcriptional regulator | cbp-2 | tm5026 | FX05026 | x0 | Deletion | Yes (Rual et al., 2004) | Not Reported | Not Reported | Not Reported |
| CTCF | CCCTC-binding factor | Component of histone acetylase and deacetylase complexes and transcriptional regulator | znf-782 | tm1265 | FX01265 | x0 | Deletion +Insertion | Not Reported | Not Reported | Not Reported | Not Reported |
| EHMT1 | Euchromatic histone lysine methyltransferase 1 | Histone methyltransferase | set-11 | ok1691 | VC1219 | x0 | Deletion | Not Reported | No (Kamath et al., 2003) | Not Reported | Not Reported |
| KDM5B | Lysine demethylase 5B | Jumonji/ARID family lysine demethylase | rbr-2 | ok2544 | RB1941 | x0 | Deletion | Not Reported | Not Reported | Not Reported | Not Reported |
| KDM6B | Lysine demethylase 6B | Jumonji/ARID family lysine demethylase | jmjd-3.1 | gk384 | VC936 | x0 | Deletion + Insertion | Not Reported | No (Kamath et al., 2003) | Not Reported | Not Reported |
| | | | | gk387 | VC912 | x0 | Deletion | | | | |
| | | | jmjd-3.2 | tm3121 | FX03121 | x0 | Deletion + Insertion | Not Reported | No (Kamath et al., 2003) | Not Reported | Not Reported |
| | | | jmjd-3.3 | tm3104 | FX03104 | x0 | Deletion | Not Reported | No (Kamath et al., 2003) | Yes (Mcdiarmid et al., 2019) | Not Reported |
| | | | jmjd-3.3 | tm3197 | FX34427 | x2 | Deletion | | | | |
| KMT2E | Lysine methylransferase 2E | Histone methyltransferase | set-26 | tm3674 | FX03674 | x0 | Deletion | Not Reported | No (Kamath et al., 2003) | Not Reported | Not Reported |
| | | | set-9 | n4949 | MT16426 | x1 | Deletion | Not Reported | No (Kamath et al., 2003) | Not Reported | Not Reported |
| KMT5B | lysine methylransferase 5B | Histone methyltransferase | set-4 | n4600 | MT14911 | x2 | Deletion | Not Reported | No (Kamath et al., 2003) | Yes (Delaney et al., 2017) | Not Reported |
| | | | | ok1481 | VC997 | x0 | Deletion | | | | |
| NSD1 | Nuclear receptor binding SET domain protein 1 | Histone methyltransferase | mes-4 | ok2326 | VC1874 | x1 | Deletion + Insertion | Not Reported | No (Carpenter et al., 2021; Kamath et al., 2003) | Not Reported | Not Reported |
| PHF2 | PHD finger protein 2 | Histone demethylase | jmjd-1.1 | tm3980 | FX17844 | x2 | Deletion | Not Reported | No (Kamath et al., 2003) | Not Reported | Not Reported |
| PHF2 | | | jmjd-1.2 | tm3713 | FX17361 | x2 | Deletion | Not Reported | No (Kamath et al., 2003) | Not Reported | No (Kleine-Kohlbrecher et al., 2010) |
| SATB1 | SATB Homeobox 1 | Chromatin remodeling scaffold | dve-1 | tm4803 | FX04803 | x0 | Deletion | Not Reported | Yes (Kamath et al., 2003) | Yes (Sheng et al., 2024) | No (Yamakazi et al., 2010) |
| SETD1A | SET domain containing 1A | Histone lysine methyltransferase | set-26 | ok952 | RB1025 | x0 | Deletion | Not Reported | No (Kamath et al., 2003) | Not Reported | Not Reported |
| SETD2 | SET domain containing 2 | Histone lysine methyltransferase | met-1 | n4337 | MT16973 | x4 | Deletion | Not Reported | Yes (Gottschalk et al., 2005) | Not Reported | Not Reported |
| | | | | tm1738 | FX01738 | x0 | Deletion | | | | |
| SETD5 | SET domain containing 5 | Histone lysine methyltransferase | set-9 | n4949 | MT16426 | x1 | Deletion | Not Reported | No (Kamath et al., 2003) | Not Reported | Not Reported |
| | | | set-24 | n4909 | MT16133 | x0 | Deletion | Not Reported | No (Kamath et al., 2003) | Yes (Mcdiarmid et al., 2019) | Not Reported |
| WAC | WW domain containing adaptor with coiled-coil | Signalling protein | wac-1.2 | ve736 | RG3236 | x0 | Deletion | Not Reported | No (Fraser et al., 2000) | Not Reported | Not Reported |
| POGZ | Pogo transposable element derived with ZNF domain | Chromatin remodeling complex protein and transcription factor | row-1 | tm556 | FX00556 | x0 | Deletion | Not Reported | No (Fraser et al., 2000) | Not Reported | Not Reported |
| ARX | Aristaless related homeobox | Homeobox transcription factor | alr-1 | ok545 | RB762 | x0 | Deletion | Not Reported | Not Reported | Yes (Topalidou et al., 2011) | Yes (Melkman and Sengupta., 2005; Tucker et al., 2005) |
| BCL11A | BCL11 Transcription Factor A | Transcription factor | bcl-11 | tm522 | FX0522 | x0 | Deletion | Not Reported | Yes (Simmer et al., 2003) | Not Reported | Not Reported |
| CIC | Capicua Transcriptional Repressor | Transcriptional repressor | gei-3 | tm4380 | FX04380 | x0 | Deletion | No (Niedworok et al., 2026) | No (Kamath et al., 2003) | Not Reported | Not Reported |
| EBF3 | EBF Transcription Factor 3 | Transcriptional activator | unc-3 | tm4776 | FX04776 | x0 | Deletion | No (Shepard et al., 2011) | No (Shepard et al., 2011) | Yes (Kim et al., 2005) | Not Reported |
| FOXG1 | Forkhead Box G1 | Transcriptional repressor | fkh-2 | ok683 | RB853 | x0 | Deletion | Not Reported | No (Kamath et al., 2003) | Not Reported | Not Reported |
| FOXP1 | Forkhead Box P1 | Transcriptional repressor | fkh-7 | gk793 | VC1646 | x0 | Deletion | Not Reported | No (Kamath et al., 2003) | Yes (Chai et al., 2026) | Not Reported |
| FOXP2 | Forkhead Box P2 | Transcriptional repressor | | | | | | | | | |

| | | | | | | | | | | | |
|---|---|---|---|---|---|---|---|---|---|---|---|
| HIVEP2 | HIVEP Zinc Finger 2 | Transcriptional activator | sma-9 | qc3 | CS67 | x1 | Point Mutation (Stop Gained) | Not Reported | Yes (Kaplan et al., 2015)/No (Kamath et al., 2003) | Not Reported | Not Reported |
| MEF2C | Myocyte Enhancer Factor 2C | Transcriptional activator | mef-2 | tm4600 | FX04600 | x0 | Deletion | Not Reported | No (Fraser et al., 2000; Rual et al., 2004 ) | Yes (van der Linden et al., 2008) | Not Reported |
| | | | | gk633 | VC1402 | x0 | Deletion | | | | |
| MYT1L | Myelin Transcription Factor 1 Like | Transcription factor | ztf-11 | ok646 | RB824 | x0 | Deletion | Not Reported | Not Reported | Not Reported | Not Reported |
| PAX5 | Paired Box 5 | Transcription factor | pax-2 | ok935 | RB1013 | x0 | Deletion | Not Reported | No (Kamath et al., 2003) | Not Reported | Not Reported |
| PHF12 | PHD Finger Protein 12 | Transcriptional repressor | athp-1 | tm4223 | FX04223 | x0 | Deletion | Not Reported | No (Gönczy et al., 2000;Kamath et al., 2003) | Not Reported | Not Reported |
| RFX3 | Regulatory Factor X3 | Transcription factor | daf-19 | tm5562 | FX05562 | x0 | Deletion | Not Reported | Not Reported | Yes (Swoboda, Adler and Thomas., 2000) | Not Reported |
| TBR1 | T-Box Brain Transcription Factor 1 | Transcriptional repressor | tbx-38 | tm581 | FX0581 | x0 | Deletion | Not Reported | No ( Gönczy et al., 2000; Kamath et al., 2003; Rual et al., 2004) | Not Reported | Not Reported |
| TBR1 | | | tbx-37 | tm314 | FX0314 | x0 | Deletion | Not Reported | No (Gönczy et al., 2000) | Not Reported | Not Reported |
| NR4A2 | Nuclear Receptor Subfamily 4 Group A Member 2 | Transcriptional regulator | nhr-6 | tm11885 | FX011885 | x0 | Deletion | Yes (Gissendanner et al., 2004, 2008) | No ( Gönczy et al., 2000; Kamath et al., 2003; Lehner et al., 2006; Rual et al., 2004) | Yes (Rossillo et al., 2020) | Not Reported |
| TCF4 | Transcription Factor 4 | Transcription factor | hlh-2 | tm1768 | FX01768 | x0 | Deletion | Not Reported | No (Rual et al., 2004) | Not Reported | Yes (Frank et al.,2003) |
| VEZF1 | Vascular Endothelial Zinc Finger 1 | Transcription factor | M03D4.4 | tm559 | FX0559 | x0 | Deletion | Not Reported | No (Kamath et al., 2003) | Yes (Mcdiarmid et al., 2019) | Not Reported |
| | | | | gk5269 | VC4183 | x0 | Deletion (Balanced) | | | | |
| ZBTB20 | Zinc Finger And BTB Domain Containing 20 | Transcription factor | ztf-6 | tm1803 | FX01803 | x0 | Deletion | Not Reported | No (Fraser et al., 2000; Rual et al., 2004 ) | Not Reported | Not Reported |
| CNOT3 | CCR4-NOT Transcription Complex Subunit 3 | mRNA deadenylase | ntl-3 | gk944863 | VC40489 | x0 | Deletion | Yes (Ceron et al., 2007) | Yes (Ceron et al., 2007) | Not Reported | Not Reported |
| ELAVL3 | ELAV Like RNA Binding Protein 3 | RNA-binding protein | exc-7 | ok370 | VC176 | x0 | Deletion | Not Reported | No (Kamath et al., 2003) | Yes (Mcdiarmid et al., 2019) | Not Reported |
| GIGYF2 | GRB10 Interacting GYF Protein 2 | Repressor of translation initiation | gyf-1 | tm4671 | FX04671 | x0 | Deletion | Not Reported | No (Kamath et al., 2003) | Not Reported | Not Reported |
| HDLBP | High Density Lipoprotein Binding Protein | RNA-binding protein | vgln-1 | ok1071 | RB1093 | x0 | Deletion | Yes (Ceron et al., 2007) | Yes (Kamath et al., 2003; Simmer et al., 2003) | Not Reported | Not Reported |
| HNRNPH2 | Heterogeneous Nuclear Ribonucleoprotein H2 | Heterogeneous nuclear ribonucleoprotein | hrpf-2 | ok1823 | VC2088 | x1 | Deletion | Not Reported | Not Reported | Not Reported | Not Reported |
| | | | hrpf-1 | tm3406 | FX03406 | x0 | Deletion | Not Reported | Not Reported | Not Reported | Not Reported |
| SON | SON DNA And RNA Binding Protein | mRNA splicing cofactor | D1037.1 | ok1746 | RB1489 | x0 | Deletion | Not Reported | No (Fraser et al., 2000) | Not Reported | Not Reported |
| CTNNB1 | Catenin Beta 1 | Transcription cofactor | bar-1 | tm8849 | FX08849 | x0 | Deletion | Yes (Kamath et al., 2003; Simmer et al., 2003) | No (Byrne et al., 2007) | Not Reported | Yes (Wu and Herman., 2006) |
| IRF2BPL | Interferon Regulatory Factor 2 Binding Protein Like | Transcription corepressor | ifbp-1 | ok1339 | VC812 | x0 | Deletion | Not Reported | No (Kamath et al., 2003) | Not Reported | Not Reported |
| RERE | Arginine-Glutamic Acid Dipeptide Repeats | Transcriptional corepressor | egl-27 | ok1670 | VC1217 | x1 | Deletion | Yes (Desai et al., 1988; Shepard et al., 2011) | Yes (Lehner et al., 2006) | Not Reported | Yes (Herman et al., 1999) |
| SIN3A | SIN3 Transcription Regulator Family Member A | Transcriptional corepressor | sin-3 | tm1276 | FX01276 | x0 | Deletion | Not Reported | No (Fraser et al., 2000; Simmer et al., 2003) | Yes (Mcdiarmid et al., 2019) | Not Reported |
| ZMYND8 | Zinc Finger MYND-Type Containing 8 | Transcriptional corepressor | bra-1 | nk1 | NU1 | x10 | Deletion | Not Reported | No (Kamath et al., 2003) | Not Reported | Not Reported |
| | | | bra-2 | ok1171 | VC869 | x1 | Deletion (Balanced) | Not Reported | No (Kamath et al., 2003) | Not Reported | Not Reported |
| DYRK1A | Dual Specificity Tyrosine Phosphorylation Regulated Kinase 1A | CTD kinase of RNAP II | mbk-1 | pk1389 | EK228 | x6 | Deletion | Not Reported | Yes (Gottschalk et al., 2005) No (Kamath et al., 2003) | Not Reported | Not Reported |
| PPP2R5D | Protein Phosphatase 2 Regulatory Subunit B'Delta | Ser/Thr Phosphatase | pptr-2 | ok1467 | RB1338 | x0 | Deletion | Yes (Rual et al., 2004) | No (Kamath et al., 2003) | Not Reported | Not Reported |

**Table S2.** ***C. elegans*** **strain information**

All ASD associated mutant strains used in this study as well as additional information including human gene name, protein function, *C. elegans* orthologue, allele, strain name, number of times outcrossed and mutation type. Phenotypic information gathered from Wormbase (https://wormbase.org/species/c_elegans#104--10), PUBMED searches for “Gene Name” AND “Sensory” OR “Dye” OR “Egg”.  Colour scheme is as follows: Green – chromatin remodellers/ histone modifiers, Orange – transcription factors, Red – RNA binding proteins, Light Blue – transcription co-factors, Purple– Kinase, Yellow – Phosphatase.

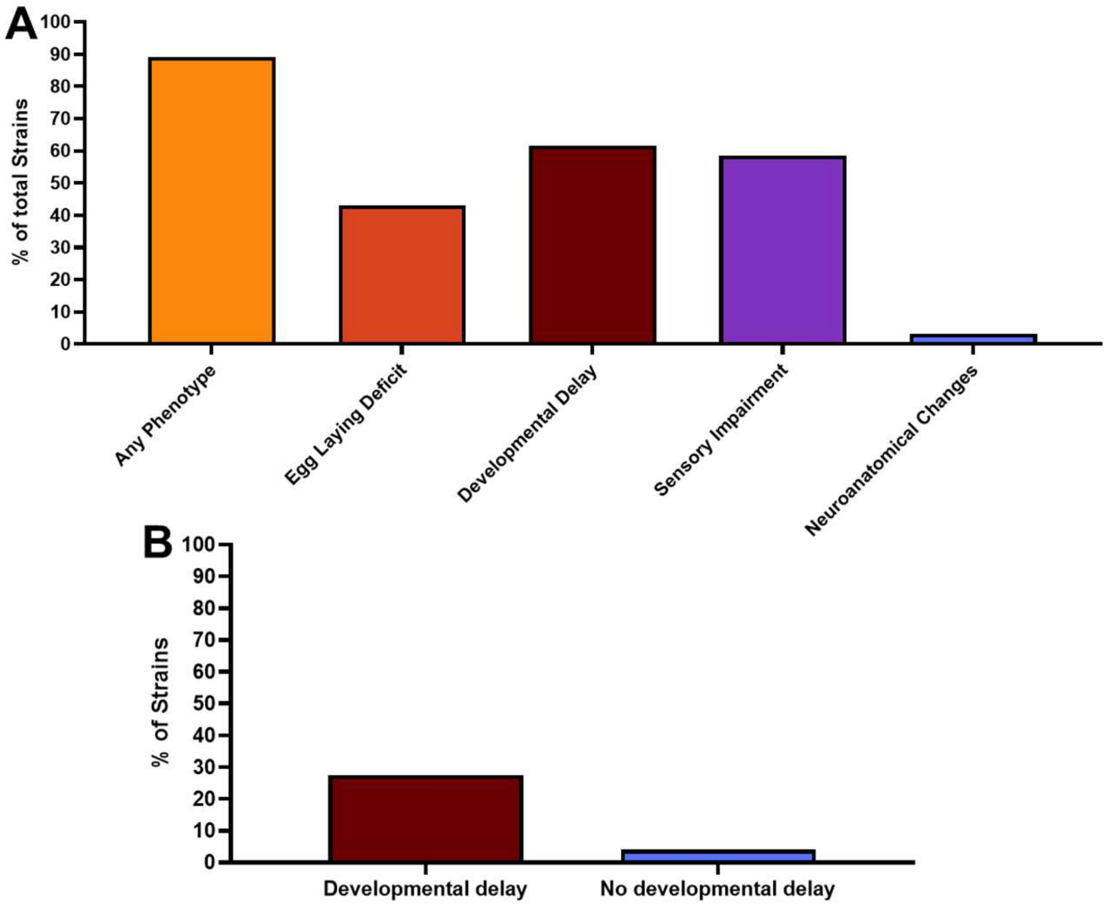

**Fig S1. Phenotypic summary of all strains investigated.**

**A.** Based on the percentage of strains with a given phenotype as outlined in figure 8. Each strain which is statistically different from N2 in any phenotype is represented under any phenotype. See methods for breakdown how each group is determined for the calculation of the % of total strains. **B.** Percentage of developmentally delayed strains with a severe sensory phenotype (See methods) compared to the percentage of strains with no developmental phenotype paired with a severe sensory phenotype.

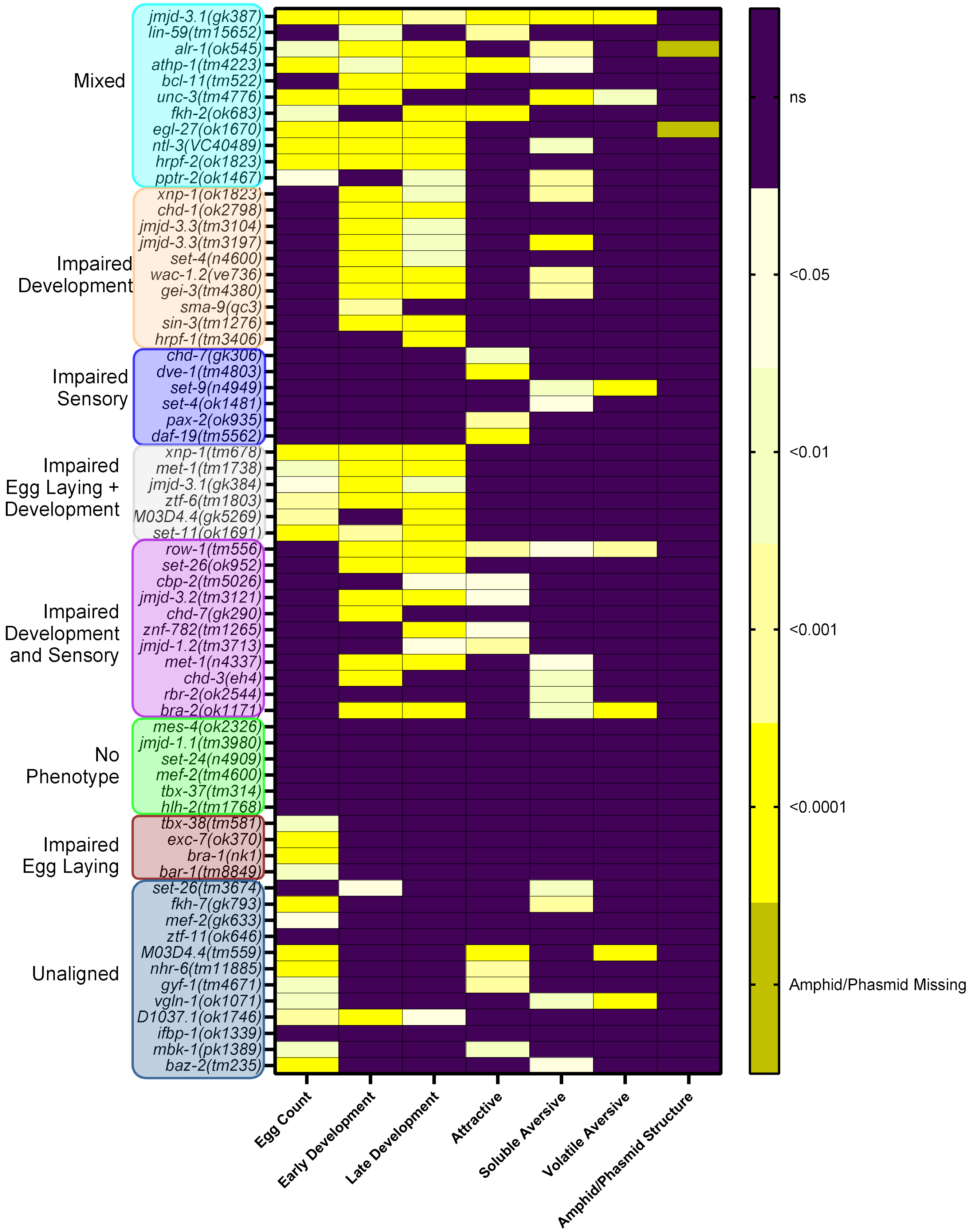

**Fig S2. Phenotypically aligned groups with corresponding significance values.**

Significance heatmap showing the seven phenotypes assessed in the experimental pipeline including, egg count, early and late development, attractive chemosensory processing, soluble aversive chemosensory processing, volatile aversive chemosensory processing and sensory amphid/phasmid structure. Strains are aligned to phenotypic groups shown on the left-hand side of the figure. Significance is defined at p≤0.05. For egg count, early and late development, attractive, soluble aversive and volatile aversive chemosensory processing one-way ANOVAs and Dunnett's multiple comparison test were performed to determine significance. For amphid/phasmid structure any deviation from the expected six amphids and two phasmids was noted as significant.
